# Hierarchal single-cell lineage tracing reveals differential fate commitment of CD8 T-cell clones in response to acute infection

**DOI:** 10.1101/2024.03.21.586160

**Authors:** Leena Abdullah, Francesco E. Emiliani, Chinmay M. Vaidya, Hannah Stuart, Fred W. Kolling, Margaret E. Ackerman, Li Song, Aaron McKenna, Yina H. Huang

**Affiliations:** Department of Microbiology and Immunology, Geisel School of Medicine at Dartmouth, Lebanon, NH 03756, USA; Department of Molecular and Systems Biology, Geisel School of Medicine at Dartmouth, Lebanon, NH 03756, USA; Thayer School of Engineering, Dartmouth College, Hanover, NH 03755, USA; Department of Biomedical Data Science, Geisel School of Medicine at Dartmouth, Lebanon, NH 03756, USA; Dartmouth Cancer Center, Lebanon, NH 03756, USA; Department of Pathology and Laboratory Medicine, Dartmouth Health, Lebanon, NH 03756, USA

**Keywords:** hierarchal lineage tracing, T-cell fate bias, endogenous CD8 T-cell response

## Abstract

Generating balanced populations of CD8 effector and memory T cells is necessary for immediate and durable immunity to infections and cancer. Yet, a definitive understanding of CD8 differentiation remains unclear. We used CARLIN, a processive lineage recording mouse model with single-cell RNA-seq and TCR-seq to track endogenous antigen-specific CD8 T cells during acute viral infection. We identified a diverse repertoire of expanded T-cell clones represented by seven transcriptional states. TCR enrichment analysis revealed differential memory- or effector-fate biases within clonal populations. Shared Vb segments and amino acid motifs were found within biased categories despite high TCR diversity. Using single-cell CARLIN barcode-seq we tracked multi-generational clones and found that unlike unbiased or memory-biased clones, which stably retain their fate profiles, effector-biased clones could adopt memory- or effector-bias within subclones. Collectively, our study demonstrates that a heterogenous T-cell repertoire specific for a shared antigen is composed of clones with distinct TCR-intrinsic fate-biases.

## Introduction

In response to viral infection, naïve CD8 T cells undergo activation, clonal expansion, and differentiation into effector cells with variable effector and survival properties. Among these effector cells are short-lived effector cells (SLECs), which express high levels of cytotoxic proteins like granzymes, terminal differentiation markers including Killer cell lectin-like receptor G1 (KLRG1), and transcription factors such as Zeb2 that repress memory differentiation ^1,2^. SLECs kill virally infected cells and then undergo programmed cell death following viral clearance. In contrast, memory precursor effector cells (MPECs) have low cytotoxic capacity but high expression of survival markers including IL-7R ^3^. MPECs persist post-viral clearance to become long-lived memory cells that are poised to establish a rapid, robust response upon antigen re-encounter. Overall, the ability of CD8 T cells to form a heterogeneous pool of functionally distinct effector cells is a core feature of adaptive immunity. Understanding how effector cell heterogeneity gives rise to a balanced proportion of effector and memory populations remains a fundamentally important question that may inform the development of better vaccines and T-cell-based therapies against infections and malignancies.

While SLECs and MPECs have been extensively studied in the context of acute viral infection, the differentiation path taken by antigen-specific T cells remains unclear. Multiple models have been proposed to explain how MPECs and SLECs arise from naïve CD8 T cells ^4–7^. Among these models are the asymmetric division model, the linear model, and the progressive (or decreasing potential) model of CD8 T-cell differentiation. The asymmetric model proposes that a newly activated naive CD8 T cell unequally distributes fate-determining transcription factors (T-bet), epigenetic regulators (Ezh2), surface proteins (CD8), and signaling components (mTOR) during the first cell division to generate effector- and memory-biased daughter cells ^8–11^. In contrast, the linear model proposes that naive T cells first differentiate into early effector CD8 T cells that can then give rise to either MPECs or SLECs ^12^. In this model, memory cells do not originate directly from naive cells but from effector cells post-viral clearance ^13,14^. Finally, the progressive model proposes that T-cell differentiation depends on T-cell receptor (TCR) signal strength, antigen affinity and accumulated signal duration. As TCR affinity and interactions with antigen-presenting cells increase, T cells lose stemness and gain effector function ^1,15^.

Additional models of CD8 T-cell differentiation have recently emerged. KLRG1 expression is associated with terminal differentiation and apoptotic cell death of SLECs. Yet, studies have shown that viral-specific KLRG1+ CD8 T cells are also present at memory time points ^16,17^. These KLRG1-expressing memory cells are called long-lived effector cells (LLEC) and provide robust protection against multiple acute infections ^16,17^. Another study showed that effector cells can lose KLRG1 expression to give rise to multiple memory CD8 T-cell subsets ^18^. Such exKLRG1 memory cells are protective in infection and tumor models ^18^.

Despite many advances in our understanding of CD8 T-cell differentiation, prior studies have largely examined the differentiation of TCR transgenic (TCR-tg) CD8 T cells following the adoptive transfer of T cells expressing super-physiological levels of a single antigen-specific TCR. These studies are also influenced by the number of transferred TCR-tg T cells, which can affect the expansion kinetics, phenotype, and even T-cell fate commitment *in vivo* ^19,20^. In comparison, only ∼100-1000 endogenous CD8 T cells are reactive to a given antigen in wild-type mice and likely express unique TCRs whose initial frequency and antigen affinity may contribute to their differentiation path ^21,22^.

Recently, the availability of T-cell tracking technologies like single-cell TCR sequencing (scTCR-seq) and T-cell static barcode labeling have enabled CD8 T-cell fate mapping ^23–26^. While these techniques are informative of the fate commitment of individual T-cell clones, they are blind to temporal and hierarchal relationships within clones. In our study, we go beyond clonal tracking of TCR-tg CD8 T cells by using unbiased approaches to fully understand the differentiation of endogenous CD8 T cells in response to acute viral infection. We use the CRISPR array repair lineage tracing (CARLIN) mouse line, which relies on CRISPR/Cas9 activity to make successive irreversible scars in a DNA barcode array as cells differentiate ^27^. The accumulation and inheritance of DNA scars within the CARLIN array are informative of the hierarchal relationships between the cells. By combining scRNA-seq and scTCR-seq with scCARLIN-barcode-seq, we have performed hierarchal fate mapping of the endogenous CD8 T-cell repertoire responding to a virally-associated antigen in its natural environment.

## Results

### A heterogeneous pool of CD8 T cells contributes to the early response to acute viral infection

To understand the lineage commitment of endogenous CD8 T cells we used the CARLIN mouse model to simultaneously capture cell fate and cell lineage history (Figure 1A)^27^. The CARLIN lineage recorder is comprised of 10 guide RNAs (gRNAs) expressed constitutively under individual U6 promoters. Each gRNA has a matching target site in a 276-bp barcode array, which is integrated into the 3’UTR of a GFP reporter gene that is ubiquitously expressed under a CAG promoter. The barcode array is targeted by doxycycline-inducible Cas9 expression. Both the CARLIN gRNAs and barcode array are expressed from the Collagen type 1 alpha 1 gene (Col1a1) locus. The reverse tetracycline transactivator (M2-rtTA) that promotes Cas9 expression in the presence of doxycycline is constitutively expressed from the Rosa26 locus. CARLIN lineage recording relies on the ability of Cas9 to make dsDNA breaks in the barcode array, which when repaired by non-homologous end-joining, result in unique deletions or insertions in the array. Subsequent rounds of Cas9 activity results in additional indels which are informative of a cells lineage history. Edited CARLIN barcode sequences, TCR sequences, and transcriptional profiles determined by scRNA-seq can then be used to understand hierarchal relationships, TCR identity, and cell states among isolated T cells.

**Figure 1.**
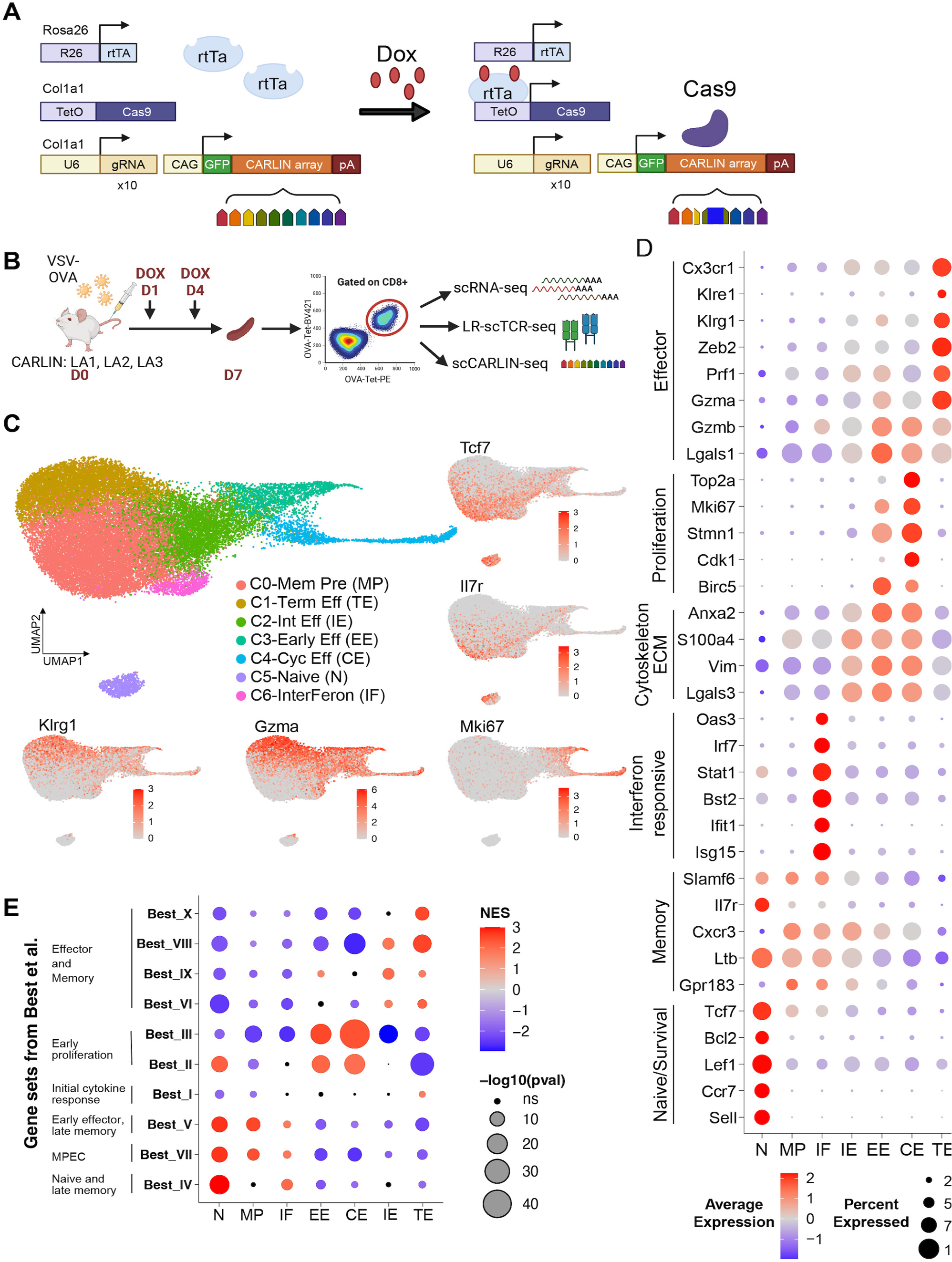
Endogenous OVA-specific CD8 T cells exhibit a spectrum of cell states at the peak of the immune response to acute VSV-OVA infection. **A.** Schematic of the lineage recorder CARLIN mouse line expressing the reverse tetracycline-controlled transactivator (rtTA) from the Rosa26 (R26) locus. Doxycycline-inducible Cas9 expression and ten U6 promoter-controlled gRNAs and CAG promoter-controlled CARLIN-barcode array within the 3’ UTR of GFP are expressed from the Col1a1 locus. The edits in the CARLIN-barcode are deletions (represented by the missing regions of the barcode) or insertions (represented by the blue blocks). **B.** Schematic of the experimental layout. Dual OVA-Tetramer+ CD8 T cells were sorted from the spleens of three VSV-OVA infected mice (LA1, LA2, LA3) on D7 post infection and analyzed by scRNA-seq, scCARLIN-barcode-seq and LR-scTCR-seq. **C.** UMAP of OVA-specific CD8 T cells from the three combined mice. Cell identities were assigned based on DEGs: Naïve-like (N), Memory precursors (MP), Early effectors (EE), Cycling effectors (CE), Intermediate effector cells (IE), Interferon responsive (IF) and Terminal effector/SLECs (TE). Feature plots of select top DEGs in CD8 T-cell clusters are also shown. **D.** Dot plot of select top DEGs in CD8 T-cell clusters. **E.** Dot plot summarizing the GSEA of gene clusters identified in Best *et al.,* which represent groups of dynamically changing genes in CD8 T cells responding to infection. The color of the dots represents normalized enrichment score (NES), and the size of the dots indicates - log_10_(adjusted p-value) of the enrichment score. ns=non-significant, ECM=extracellular matrix. See also Figure S1.

To investigate CD8 T-cell responses in an acute viral infection model, we infected CARLIN mice with Vesicular Stomatitis Virus engineered to express the model antigen ovalbumin (VSV-OVA) (Figure 1B). Two doses of doxycycline were given during the first six days post-infection to induce Cas9 expression for dynamic lineage tracing. On day 7 at the peak of the T-cell response, we sorted OVA-specific endogenous CD8 T cells using K_b_-OVA Tetramers labeled with two different fluorophores from the spleens of three infected mice (LA1, LA2, LA3) and performed scRNA-seq, long read (LR)-scTCR-seq, and scCARLIN-barcode-seq (Figure 1B, Figure S1A). We used the Seurat pipeline in R for transcriptional profiling of CD8 T cells^28^. Differential gene expression analysis followed by cell clustering using the two-dimensional Uniform Manifold Approximation and Projection (UMAP) divided the 28,875 CD8 T cells into seven distinct clusters (Figure 1C). Cells from each replicate sample was comparably represented across the Seurat clusters (Figure S1B). Based on the top differentially expressed genes (DEGs), these seven clusters represent three effector populations (C1, C3, C4), three naïve/memory populations (C0, C5, C6), and one intermediate effector population (C2) (Figure 1C-1E).

Effector T-cell clusters were enriched for transcripts associated with cytotoxic function of CD8 T cells (*Gzma, Gzmb, Prf1*) (Figure 1C-1D). Among effector clusters, C1 corresponded to terminal effector cells/SLECs (TE) as it had the highest expression of effector genes as well as markers associated with terminal differentiation of effector CD8 T cells including *Klrg1* and *Zeb2* (Figure 1C-1D). The inhibitory receptor, KLRG1, is a terminal differentiation marker of SLECs during acute infection^1^. Together with T-bet, the Zeb2 transcription factor is required to promote SLEC formation by repressing the expression of memory fate genes and upregulating effector fate genes ^2,29^. C3 and C4 were both proliferative effector populations as both had high expression of cell cycle markers (*Mki67, Birc5, Top2a*) (Figure 1C-1D). C3 cells were mainly in G2/M phase, while C4 cells were in S phase, as indicated by their Seurat cell cycle score (Figure S1C). Given the immediate proliferative burst experienced by CD8 T cells early during infection and the lower expression of SLEC markers (*Klrg1, Zeb2*) compared to TE cluster, we labeled C3 as an early effector population (EE) and C4 as a cycling effector population (CE).

Among memory clusters, C0 represented memory precursor cells (MP) as it expressed known memory markers that are important for differentiation, survival, and stemness of memory CD8 T cells (*Tcf7*, *Il7r*, *Cxcr3, Slamf6*) (Figure 1C-1D) ^3,30,31^. Apart from the MP cluster, we also found two unique populations: a naïve-like cluster (C5 or N) and an interferon-responsive memory cluster (C6 or IF). Cluster N expressed genes associated with naïve and central memory CD8 T cells (*Sell*, *Il7r*, *Ccr7, Tcf7*). Cluster IF expressed high levels of interferon-stimulated anti-viral genes (*Isg15, Bst2, Irf7, Ifit1*) and clustered closely with MP cells (Figure 1C-1D). Lastly, intermediate effector (IE) cluster C2 had low expression of memory and effector markers (*Ltb*, *Gpr183*, *Prf1*, *Lgals1*) but high expression of genes related to cytoskeleton and extracellular matrix (Figure 1D).

For each cluster, we next calculated enrichment of gene groups identified by Best *et al.* to associate clusters with various stages of CD8 T-cell differentiation in response to infection (Figure 1E) ^32^. Gene set enrichment analysis (GSEA) showed that EE and CE clusters were enriched for gene sets associated with rapid proliferation of early effector CD8 T cells (Best gene sets II, III). The IE cluster was enriched for genes expressed in effector and memory CD8 T cells (Best gene sets VI, IX, VIII). Finally, the three memory clusters (N, MP and IF) were significantly enriched for gene clusters expressed in naïve, late memory, or memory precursor cells (Best gene sets IV, V, VII). Overall, transcriptional analysis revealed a spectrum of antigen-specific effector and memory CD8 T-cell states that were present 7 days post-infection.

### Population-level RNA velocity analysis reveals early effector CD8 T cells as the parent to all effector clusters

Next, we used the RNA velocity tool scVelo to infer the differentiation trajectory of antigen-specific CD8 T-cell populations ^33,34^. RNA velocity calculates the ratio of unspliced to spliced mRNA and does not require the root cluster (precursor) to be pre-selected for analysis (Figure 2A). Using scVelo we were able to track transcriptional kinetics of DEGs without the need for sampling viral-responsive CD8 T cells from multiple mice at different time points. Initial quality checks revealed consistent proportions of total unspliced and spliced mRNA counts across the Seurat clusters (Figure S2A). scVelo identified cells in the EE and CE clusters as the root states and the TE and some IE cluster cells as the terminal states (Figure 2B). The RNA velocity trajectory predicted with high confidence that EE cells give rise to CE cells and pass through the IE cell state to differentiate into TE cells (Figure 2C, S2B). The direction of the trajectory was supported by Cytopath, a simulation-based velocity inference tool (Figure 2D) ^35^. *Zeb2* was among the top genes with differential velocity across clusters and likely informed trajectory directionality. The unspliced to spliced count ratio for *Zeb2* revealed that it was actively transcribed in the TE cluster but only modestly transcribed in EE or IE cells (Figure 2E). In contrast, RNA velocity analysis was not informative in predicting a path for MP differentiation. Nevertheless, these combined analyses suggested that early effector cells differentiate into all effector clusters (CE, IE, TE) and that they differentiate into TE cells through an intermediate effector population. This projected trajectory is supportive of the linear differentiation model, which posits that naïve CD8 T cells differentiate into early effector cells that then give rise to more differentiated effector CD8 T cells.

**Figure 2.**
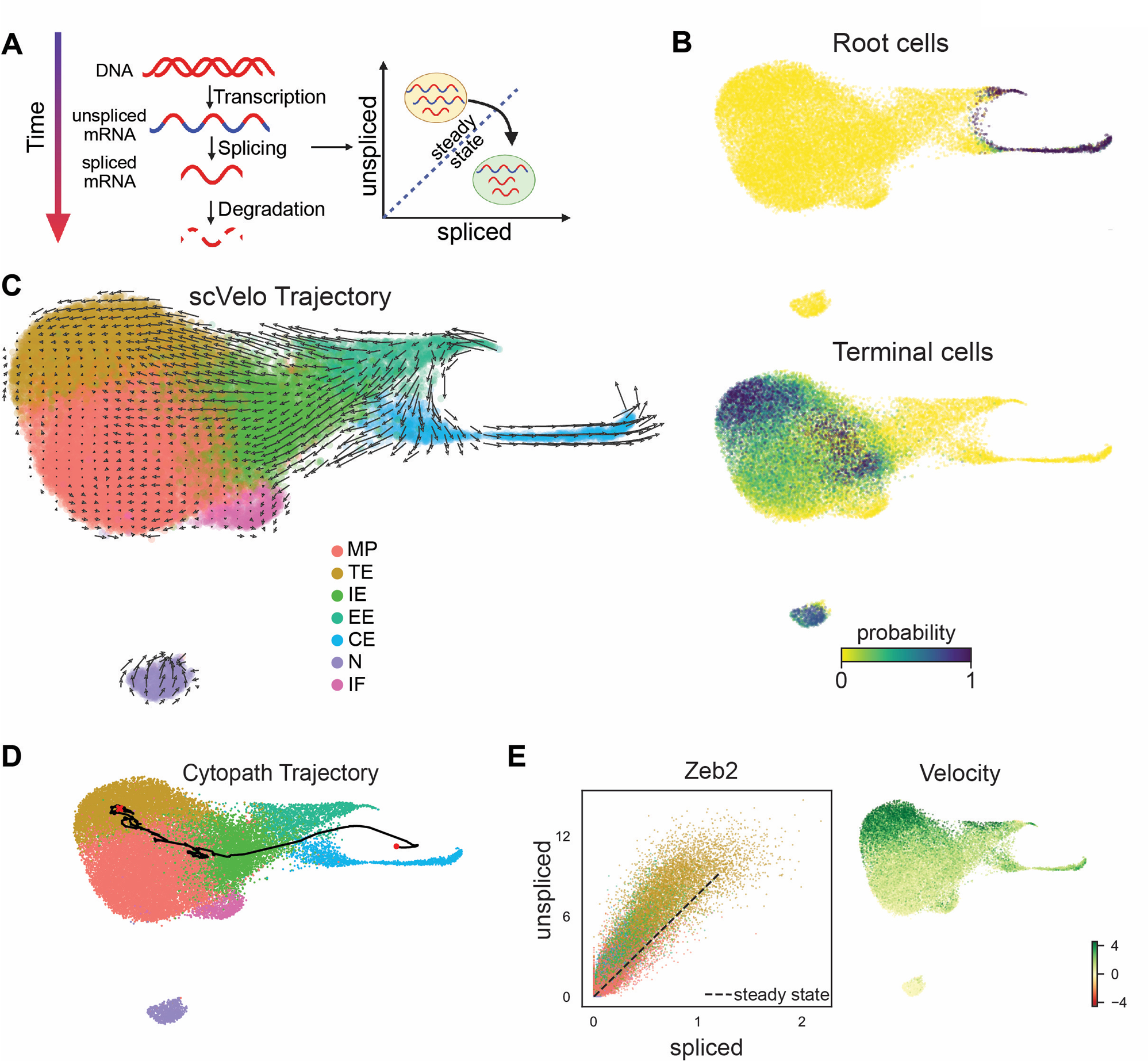
RNA Velocity reveals early effectors as the progenitors to all effector subsets. **A.** Schematic of how RNA Velocity is calculated by modeling transcriptional kinetics: rate of transcription, splicing, and degradation, which control the observed spliced to unspliced transcript ratios. **B.** Root cell and terminal cell probabilities are displayed as determined by scVelo. Velocity field projected onto the UMAP embedding as determined by **C.** scVelo and **D.** Cytopath. For scVelo, the direction of the arrows marks the direction of differentiation, and the length of the arrow indicates the rate of differentiation (velocity). For Cytopath, the red circle represents the start, and the red cross represents the end of the inferred trajectory. **E.** Scatter plot of *Zeb2* spliced and unspliced mRNA counts (left) and phase portrait of its velocity (right). The black dotted line marks the steady-state and gene velocity per cell is a measure of deviation from the steady state where positive velocity means gene induction and negative velocity means gene repression. See also Figure S2.

### Nanopore long-read sequencing uncovers numerous CD8 TCR clones that respond to acute VSV-OVA infection

Our analysis thus far has focused on the phenotype and projected fate trajectory of CD8 T cells at the population level. To investigate the role of the TCR in the fate commitment of CD8 T cells and to track individual clones, we sequenced the TCRs of the transcriptionally profiled cells. To capture the full-length receptor sequence, we enriched for V(D)J sequences and then conducted long-read Oxford Nanopore sequencing as previously described by Singh et al. in their Repertoire and Gene Expression by Sequencing (RAGE-seq) pipeline (Figure 3A) ^36^. We captured paired full-length TCR alpha and beta chain sequences of ∼85% cells in replicates LA2 and LA3 but only 17% of cells in LA1 (Figure 3B). Despite the low recovery of TCR alpha sequences from LA1, TCR beta chains were recovered for 72% of cells. For further analysis, we defined a TCR clone to comprise at least two cells that share CDR3alpha (CDR3a) and CDR3beta (CDR3b) sequences. With this criterion, we recovered 92 clones from LA1, 277 clones from LA2, and 358 clones from LA3 (Figure 3C).

**Figure 3.**
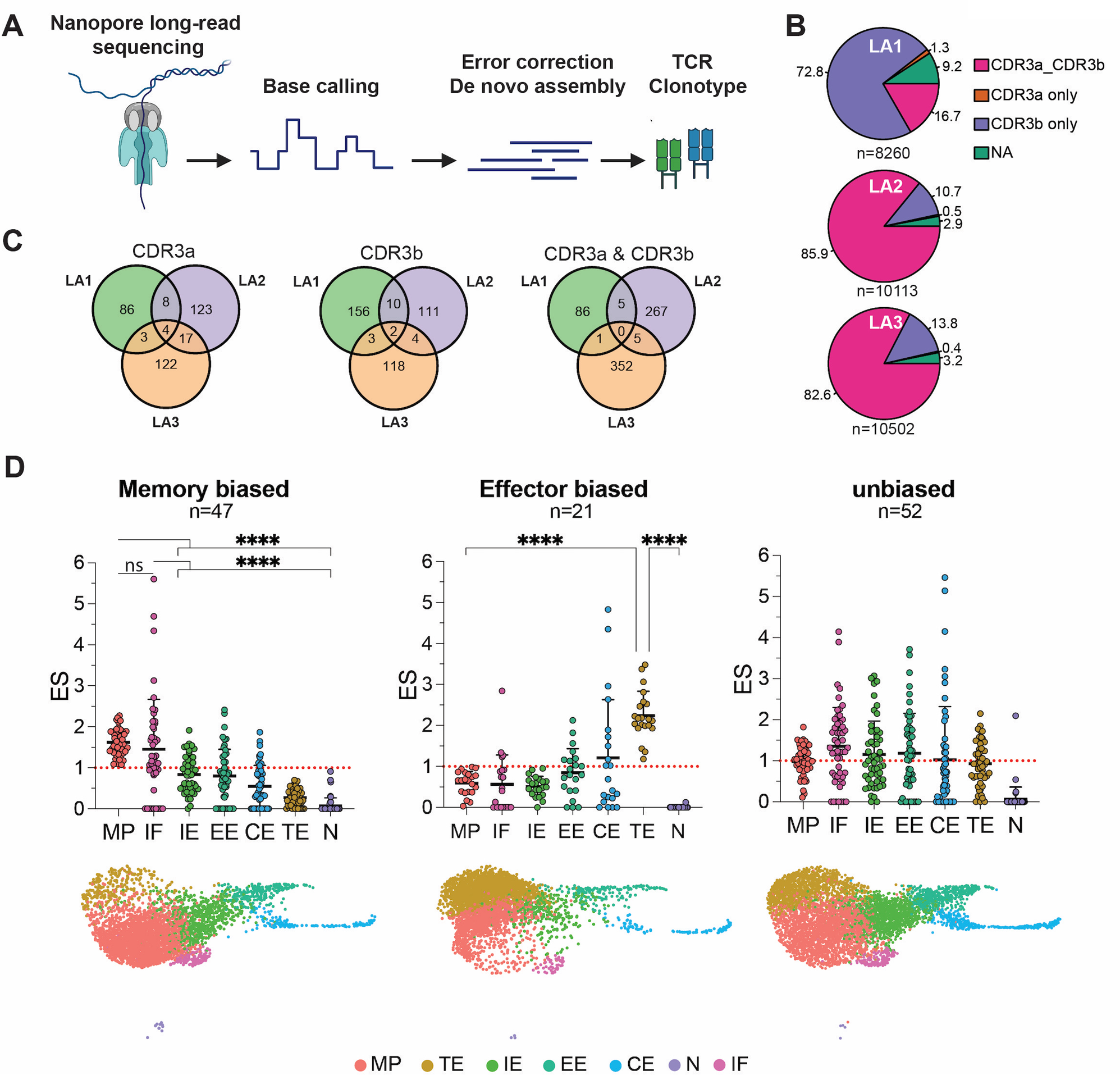
Long-read TCR sequencing reveals multiple VSV-OVA specific T-cell clones with differential fate biases. **A.** Schematic of Nanopore-based long-read TCR-seq pipeline**. B.** Pie charts depicting the percentage recovery of CDR3a and CDR3b sequences across samples. n represents the total number of T cells in each sample. **C.** Venn diagrams showing the overlap of CDR3a chains, CDR3b chains and paired CDR3a and CDR3b clones across samples. **D.** Enrichment score of the biased TCR clones in the Seurat clusters (Top). The red dotted line represents ES=1 (no enrichment). Data are individual clones (circles) and mean (bar) +SD. Statistical analysis was performed using one-way ANOVA and Tukey’s multiple comparisons test. **** P < 0.0001. Significance values for major comparisons are shown for clarity. Other significant comparisons include: IE vs TE (P <0.001), IE vs N (P <0.0001), EE vs TE (P <0.001), EE vs N (P <0.0001), CE vs N (P <0.01) for memory-biased clones, IF vs CE (P <0.05), IE vs CE (P <0.05), EE vs N (P <0.01) and CE vs N (P <0.0001) for effector biased clones and N vs all other clusters (P <0.0001) for unbiased clones. All the cells present in each biased category are displayed on the UMAP embedding (bottom). See also Figure S3.

A notable number of antigen-specific TCR clones shared CDR3a chains, CDR3b chains, and paired CDR3a and CDR3b sequences across the three replicates, despite a theoretical repertoire of ∼10^15^ TCR sequences (Figure 3C) ^37^. The captured T-cell clones varied in clonal sizes ranging from a couple to hundreds of cells. The top 20 TCR clones in LA2 and LA3 comprised ∼50% of the cells in the sample, with the most prevalent clone consisting of 1,896 cells for LA2 and 844 cells for LA3, respectively (Figure S3A). CDR3a and CDR3b VJ segment usage among the top 20 clones showed that the alpha chain was more diverse than the beta chain, with many clones in LA2 and LA3 using Vb12-1 and Vb31 segments (Figure S3B). Notably, the Vb12-1 gene segment is used by OT-1 TCR-tg T cells, which have been well characterized to recognize the OVA peptide (OVA_257-264_) used in this study to capture antigen-specific CD8 T cells ^38^. Overall, using nanopore sequencing, we recovered the full-length TCR of most of the T cells in our data and found some shared and many unique TCR sequences that respond to VSV-OVA infection.

### Expanded VSV-OVA specific TCR clones are differentially enriched for memory and effector fates

To understand the role of TCR identity on cell fate choices we calculated the enrichment of expanded TCR clones (n>10 for LA1 and n>20 for LA2 and LA3) in the Seurat clusters. Based on the enrichment scores, the TCR clones could be divided into memory-biased, effector-biased, and unbiased categories (Figure 3D). Clones that had high enrichment in the MP clusters but low enrichment in the TE cluster were labeled as memory-biased, and the ones that had the opposite preference were labeled as effector-biased. The remaining clones were assigned to the unbiased category. With this criterion, we found 47 memory-biased clones, 21 effector-biased clones, and 52 unbiased clones across the three replicates (Figure 3D). Clones in the effector-biased and memory-biased categories were significantly enriched in the cluster of interest compared to all the other clusters (Figure 3D). Notably, the memory biased clones also had significantly high enrichment in the IF cluster.

After determining the fate bias of TCR clones, we examined their VJ segment usage, amino acid properties, and position-specific amino acid composition. Differential TCR beta VJ usage was associated with fate bias (Figure 4A, top 20 clones are shown for clarity). Effector-biased clones were enriched for the Vb31 segment (6/21), and unbiased clones were enriched for the Vb12-1 segment (24/52) while memory-biased clones showed a preference for both Vb12-1 (12/47) and Vb31 (9/47) segments (Figure 4A). In contrast, TCR alpha VJ usage did not show any fate preference. CDR3 analyses revealed that effector-biased clones showed some preference for longer CDR3b sequences compared to the memory-biased and unbiased clones, but not significantly (Figure 4B). The CDR3b regions in the effector-biased clones had significantly higher average amino acid bulkiness, but significantly lower basic content and polarity compared to those in the unbiased clones (Figure 4B). However, the position-specific amino acid composition of the CDR3a and CDR3b regions in the TCR clones revealed motifs specific to each fate-category (Figure 4C, S4). ‘CASSPR’ was a dominant shared motif in twelve amino-acid long CDR3b sequences found in unbiased (8/13) and memory-biased (6/8) clones, where ‘CASS’ from Vb12-1 segment was followed by the non-germline encoded ‘PR’ sequence (Figure 4C, S4B). ‘CASSR’ was another dominant motif found in the CDR3b regions of unbiased clones (31%). Notably, ‘CASSR’ is present in the OT-1 transgenic TCR CDR3b, which can give rise to both MP and TE cells depending on the context of infection ^12,38^. Overall, the diverse pool of OVA-antigen responsive TCR clones showed differential bias for memory and effector fate but shared some intra- and inter-category features of the TCR beta chain.

**Figure 4.**
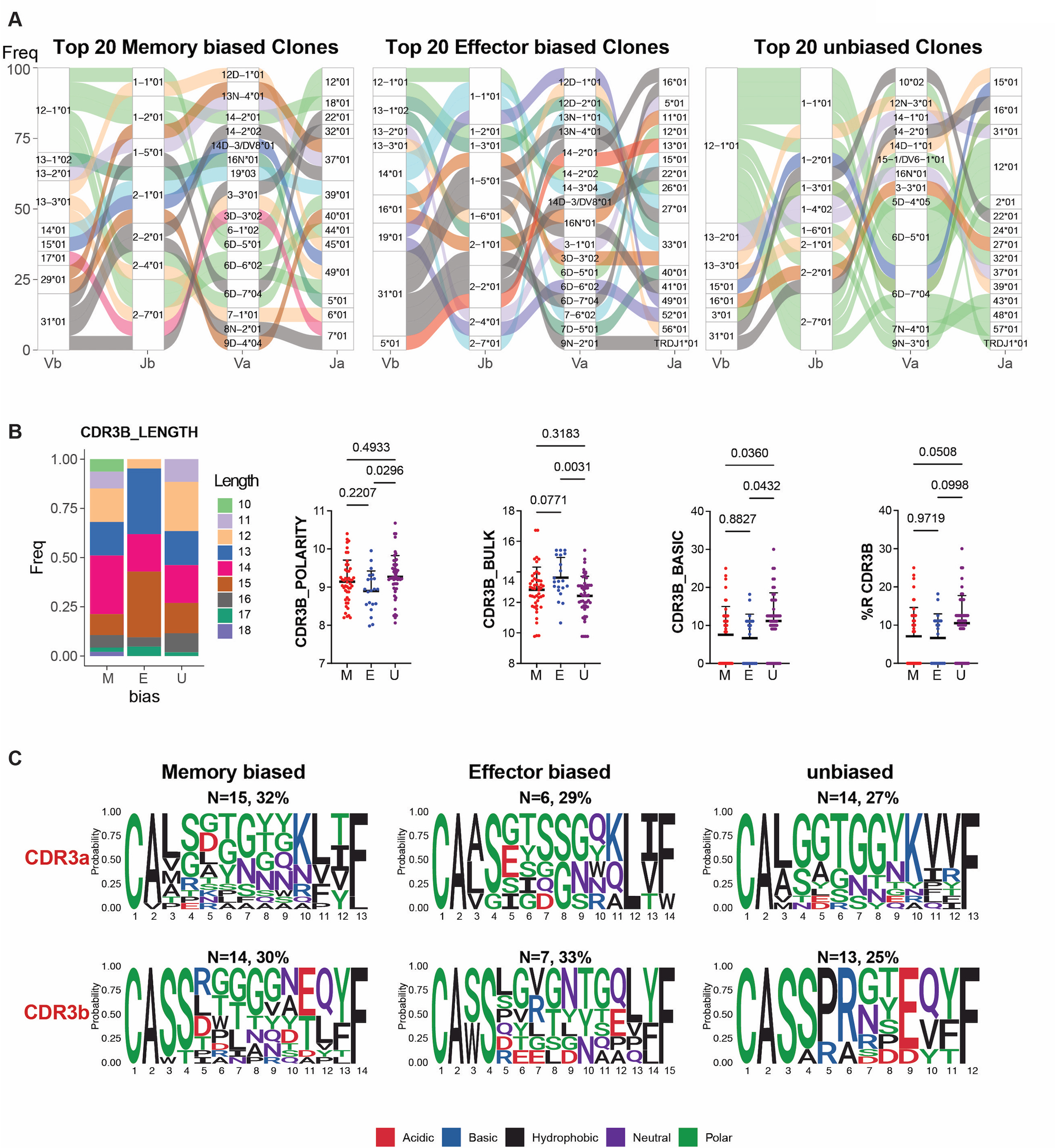
TCR sequence analysis reveals shared features within TCR fate categories. **A.** Alluvial plots showing the VJ alpha and beta usage in the top 20 clones biased towards indicated fates. The stacked bar plots represent the proportion of the TCR segments present in the top 20 biased clones. The streams connecting the stacked bar plots represent the VJ alpha beta usage for each clone. The color of the streams corresponds to specific Vb segments. Clonal size is not represented in this analysis. **B**. Length and average amino acid physiochemical properties (polarity, bulkiness, basic content, Arginine (R) content) of the CDR3b segments in the biased categories. Data are individual clones (circles) and mean (bar) +SD. Statistical analysis was conducted using one-way ANOVA and Tukey’s multiple comparisons test. **C.** Position specific amino acid composition of the dominant length CDR3a and CDR3b chains in the biased clones. The height of the amino acid letters represents their frequency at that position. The color of the letters represents the chemical property of the amino acids. Above the TCR logos, N represents the number, and the percentage represents the proportion of the biased group represented by the logo. See also Figure S4.

### CARLIN barcode edits allowed hierarchal lineage tracing of antigen-specific TCR clones

Nanopore long-read sequencing of paired TCR chains revealed fate commitment biases of the various OVA-specific CD8 T-cell clones. However, it did not uncover the differentiation pathway taken by the individual clones. In the past, such analysis required sampling of transferred TCR transgenic CD8 T cells over time from the infected host. However, the use of CARLIN mice allowed us to lineage trace endogenous OVA-specific CD8 T-cell clones collected at a single time point. Thus, we further analyzed cells that had both paired TCR and CARLIN barcode information. Since the CARLIN model uses Cas9 activity to introduce edits in the form of insertions and deletions in a ubiquitously expressed and heritable barcode, lineage trees could be built based on shared edits in the barcode for individual TCR clones (Figure 5 and 6). Within each TCR clone, cells that shared overall editing pattern in the CARLIN barcode could be grouped into subclones (Figure 5 and 6). Strikingly, for some TCR clones, accumulated successive CARLIN edits allowed us to relate subclones to inferred progenitors (Figures 5 and 6).

**Figure 5.**
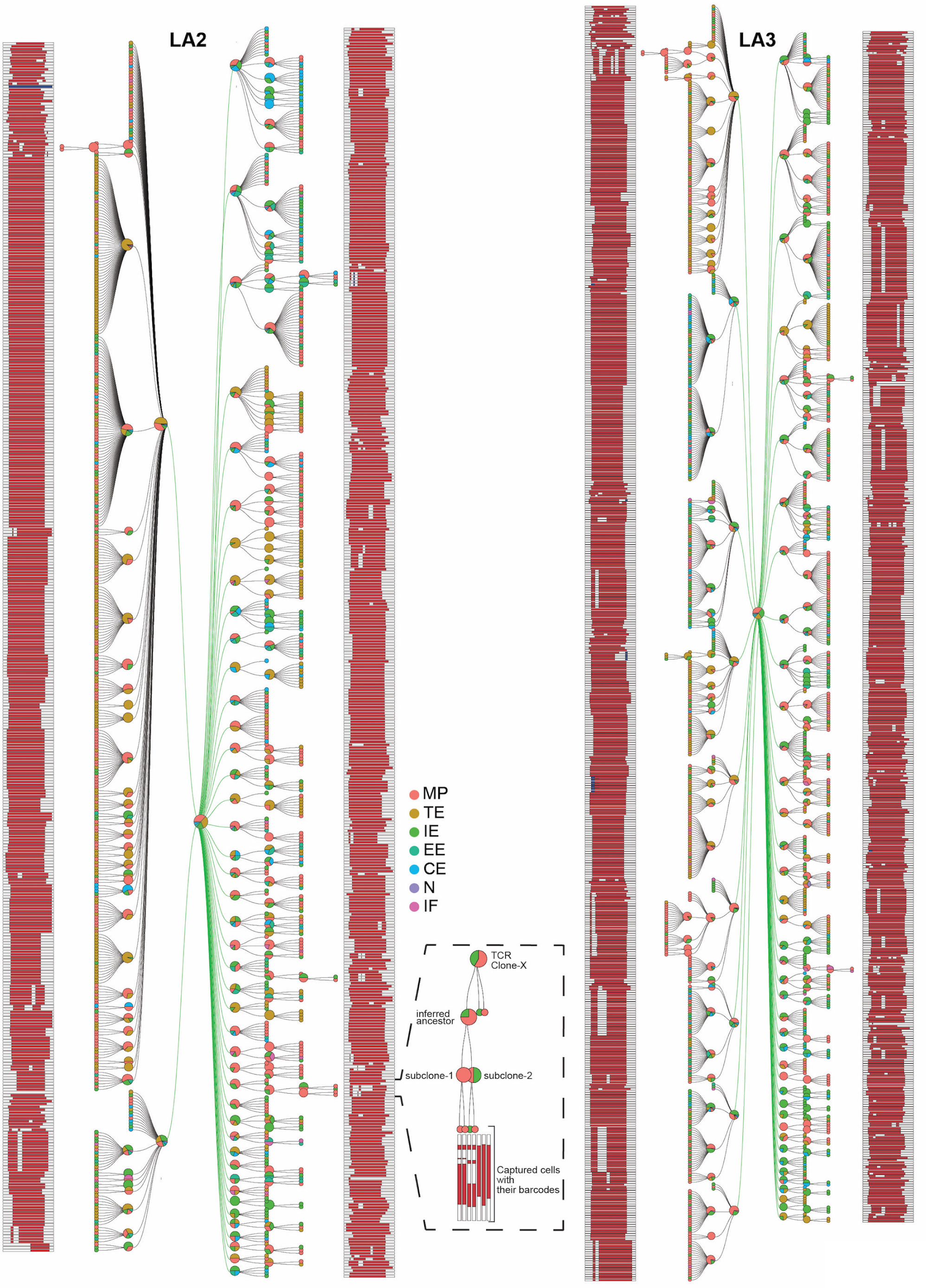
Unified Lineage Trees of TCR clones. Lineage trees for the individual TCR clones with at least three cells for which CARLIN barcode information was captured are combined into a single mega tree: LA2 (left) and LA3 (right). The individual TCR clones are connected to an arbitrary central node by green branches. Each node in the tree contains a pie chart showing the cell-type proportion of all its daughter cells. Terminal nodes are colored by their cell identity. Edited CARLIN barcodes deletions (red) and insertions (blue) are represented within white bars present on the outer edge of the trees. A sample TCR clone is highlighted in the bottom box and shows how CARLIN edits can be informative of lineage relationships.

**Figure 6.**
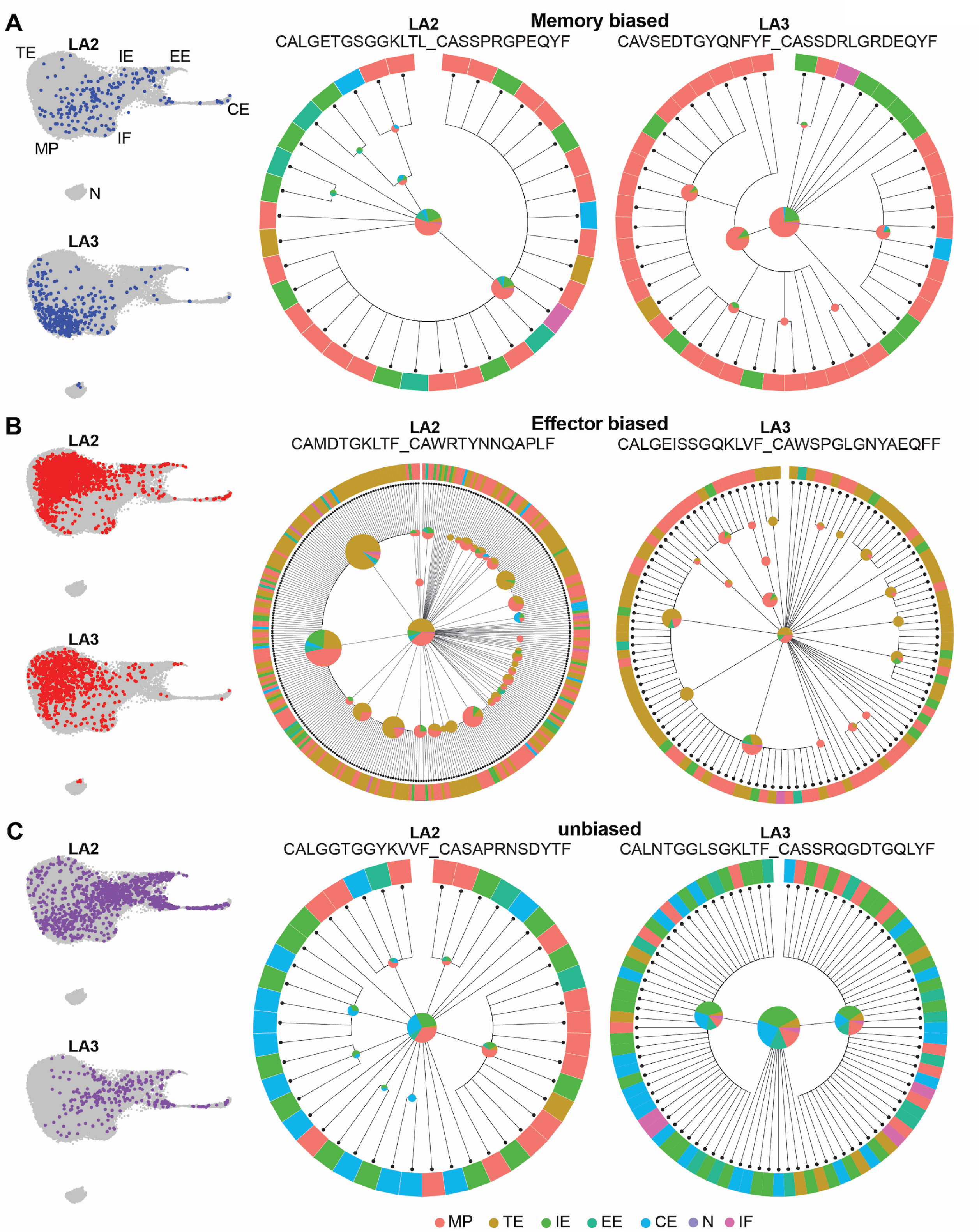
CARLIN barcode edits allow tracking of multiple generations and subclones to reveal fate biases within each TCR clone. Individual lineage trees and UMAP plots of representative TCR clones in **A.** memory-biased **B.** effector-biased and **C.** unbiased categories from replicates LA2 and LA3. Each node in the tree contains a pie chart showing the cell-type proportion of its daughter cells. The outer circle represents the cells in the TCR clones colored by their cell identity. Cdr3a_Cdr3b sequences are displayed on the top of each corresponding lineage tree. The UMAP plots show all members of a TCR clones.

We were particularly interested in comparing lineage commitment among unbiased and biased TCR clones. For this purpose, we examined phylogenetic relationships in each of the three fate-categories (Figure 6). For the memory-biased clones, similar cell fate proportions were observed within each subclone with the MP state predominating (Figure 6A). However, for the two largest effector-biased clones for which TCR and CARLIN data were recovered, we observed different subclones with a preference for effector or memory fate (Figure 6B). This unexpected result highlighted the unique value of phylogenetic analysis. Without CARLIN data, these expanded effector-biased clones would be interpreted to universally give rise to a small number of MP cells. However, the phylogenetic trees revealed that while enriched overall for effector fate, effector-biased clones could generate subclades that were generally comprised of either TE or MP cells. Finally, the unbiased clones did not show any fate preference in the subclones (Figure 6C). Thus, we speculate that the unbiased TCRs maintain their unbiased differentiation capacity; however, this needs to be confirmed by future studies. Overall, the CARLIN lineage recorder proved to be a uniquely powerful tool for tracing individual TCR clones and inferring their differentiation potential and trajectory.

## Discussion

The emergence of a balanced population of effector and memory CD8 T cells is a necessary feature of adaptive immunity. Yet, we lack a definitive understanding of how progenitor T cells progress through different transitional states to give rise to terminal effector and memory cells. In this study, using a combination of scRNA-seq, LR-scTCR-seq, and scCARLIN-barcode-seq we tracked a heterogenous pool of antigen-specific endogenous CD8 T-cell clones responding to acute viral infection. Transcriptional analysis captured multiple CD8 T-cell states ranging from early (EE) to terminal effector (TE) and memory precursor (MP) cells. Both EE and cycling effector (CE) cells were highly proliferative, less differentiated populations that may represent progenitor cells capable of replenishing the effector and memory compartments. The ability of EE and CE cells to act as precursors was supported by RNA velocity analysis, which identified them as root states with velocity trajectory projecting that EE cells transition through IE cells to give rise to TE cells. Among the genes with the most differential velocity used for predicting the overall velocity trajectory was Zeb2, which was actively transcribed in the TE cluster (high unspliced to spliced ratio). Zeb2 is a transcriptional repressor that acts downstream of T-bet to repress memory genes (*Il7r, Tcf7*) and has been shown to be essential for the formation of KLRG1-expressing terminal effector cells ^2,25,29^. Future studies to determine whether EE cells exiting the cell cycle upregulate Zeb2 expression to give rise to IE cells prior to commitment to the TE fate will be required to support the terminal differentiation sequence predicted by trajectory analysis.

In contrast to TE cells, RNA velocity failed to clearly infer the differentiation trajectory of MP cells. However, the IE population expressed both effector and memory markers. Additionally, most memory-biased clones showed some enrichment in the IE cluster, providing support that IE cells may also differentiate into MP cells. IE cells potentially represent a transitional cell state with multiple differentiation potentials. Kasmani *et al.* recently described a bifurcated CD8 T-cell differentiation model, in which an intermediate CD8 T cell population that shared top DEGs (*Vim*, *S100a4*, *S100a6*, *S100a10*, *Lsp1*, *Lgals1*, *Lgals3*) with IE cells can differentiate into either terminal effector or exhausted CD8 T cells in response to chronic viral infection ^25^. It will be important to conduct future adoptive transfer experiments with endogenous IE cells to examine their full differentiation potential.

We also uncovered a unique memory CD8 T-cell population that appeared to be Interferon Responsive (IF). IF cells had high expression of Type I interferon (IFN) related genes and memory markers. Prolonged Type I IFN signaling is associated with increased CD8 T-cell exhaustion during chronic infections ^39,40^. However, in certain acute infections Type I IFNs can act directly on CD8 T cells to promote their survival as they proliferate in response to antigen to form an optimal pool of memory cells ^41^. More recent studies have identified an important function for Type I IFN signaling in promoting the formation of tissue-resident memory (TRM) T cells. Molodtsov and Khatwani *et al.* uncovered a population of CD8 TRM cells in the lymph node that co-express *Il7r* and Type I IFN signaling genes in a mouse melanoma model ^42^. Moreover, Varese *et al.* showed that early exposure to type I interferons during primary viral infection is required for lung CD8 TRM expansion upon re-infection ^43^. Future lineage tracing studies that assess memory T cells from lymphoid and non-lymphoid tissues are required to better understand the developmental relationship between IF and MP cells and circulating versus tissue-resident memory T cells.

Transcriptional analysis also revealed a small Naïve-like (N) population. We refer to this cluster as Naïve-like because they have high expression of naïve and survival markers (*Il7r, Sell, Tcf7, Bcl2*) and the early activation marker *Cd69* (data not shown). N cells were largely absent from the expanded biased TCR clones, leading us to propose that N cells may represent T cells that are recently or non-productively activated.

We next asked whether different OVA-specific TCR clones favored distinct cell states. Due to the highly diverse nature of TCR sequences generated from random nucleotide insertions and deletions at the VJ junctions, we required an accurate long read sequencing method. Recent advances in the field enabled us to obtain full-length paired TCR alpha and beta chain sequences using Oxford Nanopore technology as described by Singh et al in their RAGE-seq study^36^. However, in the original RAGE-seq study TCR recovery was low due to high error rate in the nanopore data. Improvements in the Nanopore sequencing accuracy coupled with the two-step error correction method that we developed allowed us to recover ∼85% of paired TCR sequences from LA2 and LA3 replicates. This increase in recovery was critical for multi-omic analysis of a large proportion of the antigen-specific T cells.

Even though we captured CD8 T cells specific for a single antigen in genetically identical mice, our data uncovered a diverse pool of expanded OVA-specific CD8 T-cell clones. Previous studies have also reported high TCR diversity in response to a single antigen ^25,44^. Moreover, we discovered that different CD8 T cell clones had different fate biases. Most of the clones belonged to the memory-biased and unbiased groups. Clones in the memory-biased category were also significantly enriched in the IF cluster compared to all the effector clusters, further supporting the possibility that IF cells are a true memory precursor population. Within each fate-category, some shared sequence-based properties were evident despite high diversity in the TCR clones. The unbiased clones mainly utilized the Vb12-1 segment, and the effector-biased clones showed a preference for the Vb31 segment. It was interesting to see that the unbiased clones preferred the same Vb that is found in the OT-1 TCR that has long been used to study CD8 T-cell differentiation in viral and bacterial infection and cancer models. The OT-1 CDR3b sequence also contained ‘CASSR’, which was a dominant motif in the CDR3b sequence of the unbiased clones (∼33%) ^38^.

CARLIN lineage tracing allowed us to follow generations of the same TCR clones without the need for resampling. We could follow multiple subpopulations of TCR clones, and in some cases, also track sequential generations of subclones. scRNA-seq combined with LR-scTCR-seq revealed the fate preferences of each clone, but scCARLIN-barcode-seq was required to track the differentiation histories within each clone. For the unbiased clones, we were interested to see whether fate bifurcation happens as inferred for the overall population of CD8 T cells using RNA velocity and TCR enrichment analysis. The observed subclones from unbiased and memory-biased clones did not show any change in fate preference. However, some effector-biased clones included subclones that were dominated by either TE or MP fates. We speculate that subclones within the effector-biased category may bifurcate by continuing to integrate extrinsic environmental cues to make future cell fate choices. In contrast, intrinsic factors such as TCR sequence identity may continue to dominate the fate commitment of unbiased and memory-biased clones.

To the best of our knowledge, our study is the first to use Cas9-based hierarchal lineage tracing to track the differentiation of endogenous CD8 T cells. Much of our understanding of viral-specific CD8 T-cell differentiation has come from OT-1 T-cells and their ability to give rise to MP and TE cells depending on environmental cues ^12^. The current study links CD8 T-cell TCR sequences to their transcriptional profile. However, we are unable to elucidate the impact of extrinsic factors on the CD8 T-cell differentiation such as APC identity, tissue localization, and microenvironment ^45^. Previous lineage tracing studies transferred a single TCR-tg T cell ^46^ or a pool of TCR-tg T cells that express static barcodes ^23,24^. But these studies required removing CD8 T cells from their natural environment and restricted analysis to a single TCR. A physiological response, however, engages a relatively large repertoire of T cells. Our data suggests that TCR identity imparts intrinsic fate biases that may ensure balanced proportions of memory and effector CD8 T-cell clones, which serve specialized functions required for optimal immediate and long-term host protection.

There are multiple models for CD8 T-cell effector and memory differentiation. No single model of differentiation fits all the OVA-specific TCR clones characterized. At least for the biased clones, it appears unlikely that their fate choice followed the asymmetric model of differentiation since their TCR sequences were selectively represented in either effector or memory cells. RNA velocity combined with TCR enrichment scores support the linear model of differentiation, in which the early effector cells give rise to TE cells and possibly MP cells in a TCR-dependent manner. Lastly, we were unable to directly evaluate the progressive model of differentiation, which requires the determination of TCR avidity and interaction dynamics with antigen-presenting cells (APCs). Future improvements to lineage tracing models that would allow recording of accumulated TCR signals and improve the resolution of heritable barcode edits are needed to fully evaluate the linear and progressive differentiation pathways ^47^. Nevertheless, mechanistic analyses of how TCR bias influences the T cell repertoire responding to an antigenic insult will not only improve our fundamental understanding of the aggregated T-cell response but may also have a striking impact on the design of T-cell-based therapies against viruses and cancer.

## Author contributions

L.A. and Y.H.H. conceived the project, designed all experiments, and wrote the original manuscript. All co-authors provided feedback on the final manuscript. L.A., F.E.E., and F.W.K. performed the experiments. L.A., C.M.V., H.S., L.S., and A.M. processed and analyzed the data using new and modified pipelines. M.E.A., A.M. and Y.H.H. supervised and secured funding for the study.

## Acknowledgments

We thank Gary Ward for flow cytometry support, Eric DuFour for animal husbandry, and Shawn Musial for help with VSV-OVA infections. This work was supported in part by National Institutes of Health (NIH) grant R01-AI131975 to M.E.A and Y.H.H., DP2-GM149750 to A.M., V Foundation for Cancer Research and Pew Biomedical Scholars awards to A.M., Tom and Susan Stepp to Y.H.H, Prouty Developmental grant to A.M. and Y.H.H., NIH grants P20-GM130454, S10-OD025235 and S10-OD030242, which supports the single cell genomics core, and NIH grant P30-CA023108, which supports the Dartmouth Cancer Center’s flow cytometry and genomics cores. Some figure schematics were generated in Biorender.com.

## STAR Methods

### Lead Contact

Further information and requests for resources and reagents should be directed to and will be fulfilled by the lead contact, Yina H. Huang.

### Materials availability

This study did not generate new unique reagents.

### Mice

CARLIN (RRID:MMRRC_067061-JAX) and KH2/iCas9 (RRID:IMSR_JAX:029415) mice were purchased from The Jackson Laboratory ^27^. rtTa and Cas9 were bred out of the CARLIN strain at The Jackson Laboratory. As a result, CARLIN mice were crossed with KH2/iCas9 mice. The F1 generation that is heterozygous for the CARLIN barcode, tetO-Cas9 and rtTa locus was used in this study. All mice were housed in specific pathogen-free facilities and cared for in accordance with the guidelines of Dartmouth College. All animal studies were approved by the Institutional Animal Care and Use Committee of Dartmouth College.

### Virus and infection

Seven-week-old male CARLIN mice were infected intravenously with 1 × 10^6^ plaque-forming units (PFU) of VSV-OVA kindly provided by Dr. Pamela Rosato (Dartmouth College) ^48^.

### Doxycycline Dosage

Mice were given two ‘pulses’ of doxycycline (Sigma-Aldrich, SKU: D9891). Doxycycline was administered to the mice in drinking water (2mg/ml with 10mg/ml sucrose) and through intraperitoneal (IP) injections (2mg/200μl of 1xPBS, per mouse). Mice were kept on doxycycline water on days 1 and 2, and then days 4 and 5 post-infection, and administered doxycycline by IP injections on days 1 and 4 post-infection.

### Class I MHC OVA-Tetramer Preparation

Class I MHC H-2K(b) biotinylated monomers containing the OVA-derived peptide SIINFEKL were kindly provided by NIH Tetramer Core Facility (contract number 75N93020D00005). The tetramerization was done as per manufacturer’s instructions with PE-Streptavidin (BioLegend, Cat#405203) and BV421-Strepavidin (BioLegend, Cat#405226).

### Tissue preparation and staining

After 7 days of infection, 3 CARLIN mice were euthanized, and their spleens were harvested. The spleens were mashed in flow staining buffer (1xPBS, 2%FBS, 1mM EDTA) using the back of a 5ml syringe plunger and then passed through a 70-micron filter to get a single cell suspension. The splenocytes were treated with 1x red blood cell lysis buffer (BioLegend, Cat#420302) for 5 minutes (mins) at room temperature (RT) to deplete erythrocytes. Next, CD8 T cells were enriched in flow buffer using the EasySep™ Mouse CD8+ T-cell Isolation Kit as per manufacturer’s instructions (STEMCELL, Cat#19853). Enriched CD8 T cells were then incubated with Zombie NIR dye (BioLegend, Cat#423106) in 1xPBS for 10 mins at RT in darkness followed by a wash with excess 1xPBS. The enriched CD8+ T cells were then incubated with APC anti-mouse CD8a antibody (BioLegend, Cat#100712, RRID:AB_312751), OVA-Tetramer-BV421 and OVA-Tetramer-PE for 30 mins on ice in darkness. Stained cells were resuspended at about 5 million cells/ml concentration in flow buffer for the cell sort. Finally, live, OVA-Tetramer-BV421 and OVA-Tetramer-PE double positive CD8+ T cells were sorted in 1xPBS + 0.05% BSA on a SONY SH800S cell sorter.

### sc-RNAseq library construction and sequencing

Cell suspensions were prepared in 1xPBS + 0.05% BSA and cell concentration and viability measured on a K2 automated fluorescence cell counter with AO/PI dye (Nexcelom Bioscience). Cells were loaded on to a 10x Genomics Chip G targeting 10,000 cells/sample and processed using the 10x Genomics NextGEM 3’ v3.1 chemistry (PN-1000268). Libraries were prepared according to manufacturer’s instructions and sequenced using NextSeq2000 paired-end 100 cycle kits (Illumina, Cat#20040559), targeting 25,000 reads/cell (Read1: 28 cycles; Read2: 90 cycles; Index1: 10 cycles; Index2: 10 cycles).

### CARLIN barcode library construction and sequencing

CARLIN libraries were prepared as described in the original CARLIN study ^27^. Briefly, for CARLIN barcode amplification, 5ng of amplified cDNA from step 2.3 of the 10x 3’ v3.1 User Guide (CG000204 Rev D) was PCR amplified (10X-CARLIN_1-bio, P5-PR1; 15 cycles), enriched with kilobaseBINDER streptavidin beads (Invitrogen, Cat#60101), and further amplified (10X-CARLIN_2, P5-PR1; 15 cycles) prior to 0.8X SPRIselect cleanup. Samples were indexed using the 10x Genomics “TT” dual-index plate (PN-1000215). Sequencing was performed on a NovaSeq6000 using SP 500 cycle flow cells (Cat#20028402; Read1: 28 cycles; Read2: 300 cycles; Index1: 10 cycles; Index2: 10 cycles).

### LR-scTCR library construction and sequencing

LR-scTCR-seq libraries were prepared using a modified version of the protocol described in the original RAGE-seq paper ^36^. After preparation of 10x RNA-seq and CARLIN libraries, the remaining ∼20-30ul of cDNA from step 2.3 of the 10x 3’ v3.1 User Guide (CG000204 Rev D) was used as input into the RAGE-seq protocol. cDNA was amplified for 20 cycles using primers against the 10x TSO (AAGCAGTGGTATCAACGCAGAGT) and Partial TruSeq Read 1 sequences (CTACACGACGCTCTTCCGATCT). Then V(D)J sequences were enriched with the xGen Hybridization Capture chemistry (IDT) using a custom probe set. Probes were designed against all mouse functional V, D, J and TCR/BCR constant regions obtained from the IMGT database. Following hybridization capture, libraries were further amplified 20 cycles to have sufficient material for Nanopore sequencing. Each sample (1 per 10x library) was prepared for nanopore sequencing using the ligation sequencing kit for either the V10 (Sample LA1, SQK-LSK110) or V14 (Samples LA2 and LA3, SQK-LSK114) chemistry and sequenced using the corresponding FLO-MIN110 or FLO-MIN114 flow cells. Super accuracy base calling was performed off instrument using dorado v0.2.1 on an Nvidia A40 GPU. Sequencing yields averaged ∼2.5Gb for V10 and ∼15Gb for V14 runs.

### 10x Genomics data pre-processing

10x Genomics 3’ v3.1 data were processed using the Cell Ranger count (v6.0.1) to generate gene expression matrices for downstream processing.

### scRNA-seq data analysis

Transcriptional profiling of the captured CD8 T cells was done mainly in R using the Seurat package (v4.3.0.1) ^28^. For the 3 samples (LA1, LA2, LA3), the feature-barcode matrices from Cell Ranger were merged using the *merge()* function into a single Seurat object. Cells in which less than 1000 or greater than 6000 unique genes were detected and that expressed 7% or more transcripts corresponding to mitochondrial genes were filtered out. Also, genes that were expressed in fewer than 3 cells/sample were removed. After quality control, the count matrix comprised of 16,256 genes detected in 28,875 cells in total. The count matrix was normalized using ‘LogNormalize’ method. Then to ensure that the downstream analysis was focused on the genes with the most variable expression across cells the top 2,000 most variable genes were identified using the *FindVariableFeatures()* function followed by a linear transformation or scaling, which prevents bias towards highly expressed genes. Next, Principal Component Analysis (PCA) dimension reduction was performed on the scaled data using the top 2,000 most variable genes. Visualization of PC_1 vs PC_2 plot revealed that the samples were not well integrated. Thus, the Harmony (v0.1.1) algorithm was used to correct for dataset biased cell clustering ^49^. The cells were then clustered using the first 15 harmony-corrected PCs and a cluster resolution of 0.25 (determined by *clustree()* function). The clusters were visualized in two-dimensional space with the dimensional reduction technique Uniform Manifold Approximation and Projection (UMAP) using the first 15 harmony-corrected PCs. Differential gene expression analysis was performed using the *FindMarkers()* function for each clusters versus all the other cells using the non-parametric Wilcoxon Rank Sum test. Top DEGs in each cluster were used to identify the CD8 T-cell states/type they represent.

### Gene Set Enrichment Analysis (GSEA)

GSEA was performed using fgsea package (v1.22.0) in R (https://bioconductor.org/packages/release/bioc/html/fgsea.html). For each cluster, DEGs were determined using *FindMarkers()* function and then ranked by their average log_2_ fold change. Gene signatures corresponding to various stages of CD8 T-cell differentiation in response to infection were obtained from Best et al. 2013 ^32^.

### RNA velocity trajectory analysis

RNA velocity models transcription kinetics using the unspliced and spliced transcripts of genes. The BAM files from cell ranger output were fed into the velocyto.py pipeline (v0.17.17) ^34^ to generate .loom files that store spliced and unspliced count matrices for each sample. RNA velocity trajectory analysis was performed using scVelo tool (v0.2.5) ^33,34^. First, the Seurat object was converted into an anndata object (https://anndata.readthedocs.io/en/latest/) using modified code from https://smorabit.github.io/tutorials/8_velocyto/ so that it was compatible with the scVelo pipeline in python. The anndata object and the loom files for the 3 samples were merged into a single object for velocity estimations. The data matrix was filtered and normalized and genes that expressed fewer than 20 counts of both spliced and unspliced mRNA were removed. First and second order moments were computed for velocity estimation among nearest neighbors using the first 15 harmony-corrected PCs and 20 neighbors with the umap method. Velocity was calculated using the ‘stochastic’ model. The root cell and terminal cell probabilities for each cell were computed using the *scv.tl.terminal_states()* function. The calculated velocities were projected onto the UMAP generated from the Seurat pipeline. Genes with differential velocity expression across Seurat clusters were identified and ranked using *scv.tl.rank_velocity_genes()* function.

### Cytopath

Cytopath (v0.1.9) is a simulation-based method to infer differentiation trajectories ^35^. It takes the cell-to-cell transition matrix (excluding self-transitions), terminal states and initial states data from scVelo as input. It then simulates differentiation from the initial clusters to the terminal clusters and generates a trajectory. The cytopath-inferred trajectory was projected onto the UMAP generated from the Seurat pipeline. The terminal effector (TE) cluster was used as the end cluster for the trajectory inference as it had the highest average terminal state probability.

### LR-scTCR-seq data analysis

First, raw .fastq files were split into temporary sub-files for faster processing time via the command seqkit split2 ^50^ and imported into R. Levenshtein distances were computed for forward (Partial TruSeq Read 1) and reverse (TSO, PN-3000228) Illumina adapters, termed “r1” and “polydt”, respectively. The reverse complement sequences were labeled as “r1_rev” and “polydt_rev”. Putative barcodes were defined as the 16-nucleotide sequences located adjacent to the “r1” and “r1_rev” sequences. These putative barcodes were cross-checked against the Cell Ranger whitelist (“3M-february-2018.txt.gz”). Following this check, two sets of error correction were then applied.

In the first error correction stage, concatenated sequences were identified and corrected. These were defined as sequences having perfect matches to at least three of the four following primers: “r1”, “r1_rev”, “poly_dt”, and “polydt_rev”. Sequences with adjacent adapters in the middle were split and labeled, and each resulting sequence had its barcode extracted individually. The second error correction focused on barcodes not found in the whitelist. First, a contingency table of incorrect barcodes was constructed. This table ranked each unique incorrect barcode by its frequency of appearance. The top 10,000 incorrect barcodes, based on the number of associated reads, were selected for error correction. Each incorrect barcode was processed as follows: all barcodes with an edit distance of one (indicating a single base mutation) were generated (altered barcodes). This list of altered barcodes was screened first against all correct barcodes appearing in the dataset, and then against the whitelist. Altered barcodes matching a “correct” whitelist barcode were assigned to the corresponding incorrect barcode. Among these corrected barcode reads only those that had exactly one matching ‘correct’ barcode were kept and the rest were discarded.

The entire dataset was subsequently divided into individual .fastq files per cell. Each file, and its corresponding set of sequences, underwent error correction using Canu ^51^ with the command *canu -correct*, and the parameters *stoponLowCoverage = 0.1, minInputCoverage = 0.1, genomesize = 15k, minReadLength = 100, minOverlapLength = 50,* and *correctedErrorRate* = 0.4. Post-Canu processing, every error-corrected read from a cell was subjected to IgBLAST (v1.20.0), with the organism set to mouse and using the IMGT germline libraries for V, D, and J sequences. Following IgBLAST, all reads were reimported into R on a per-cell basis. Cells containing more than three full-length, high-quality VJ reads were selected for further analysis. For each cell, the full-length receptor with the most reads having identical VJ and CDR3 annotations was assigned. Each individual TCRa and TCRb assignment was then appended to the metadata of the processed Seurat object for subsequent visualization and analysis.

To assess the reliability of this pipeline’s assignment with gold-standard TCRa/TCRb pipelines, we compared the resulting VJ and CDR3 assignments to those from TRUST4(v1.0.6) ^52^, which was used independently to annotate receptor identity on the scRNA-seq datasets from the three samples. Cells that passed Seurat QC and had both TRUST4 and LR-scTCR-seq assignments were compared based on VJ and CDR3 identity. The TRUST4 and LR-scTCR-seq assignments matched in more than 95% of the cells.

### Enrichment Score calculation and biased clone assignment

To determine if certain TCR clones were enriched in a particular Seurat cluster an ‘Enrichment Score’ (ES) was calculated. In a sample, ES for a given TCR clone 1 in a cluster X is calculated as: 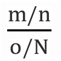 where:

N=total cells in the sample

n=total cells in TCR clone 1

m=number of cells in TCR clone 1 that belong to cluster X

o=number of cells in the sample that belong to cluster X

Cells with no TCR information were removed prior to this analysis. ES was determined for only expanded T-cell clones (n>10 for LA1 and n>20 for LA2/LA3) in all Seurat clusters. An ES of 1 indicates no enrichment, >1 indicates enrichment and <1 indicates depletion.

To assign a clone to the memory-biased group it had to satisfy three conditions:

1. The clone should have ES>1 in the MP cluster
2. The clone should have ES<1 in the TE cluster
3. 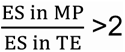

The opposite criteria had to be satisfied for clones to be assigned to the effector biased group. All the remaining TCR clones were assigned to the unbiased category.

### CDR3 amino acid property analysis

The average physiochemical properties of the CDR3b segments (polarity, bulkiness, basic content) were determined by Alakazam package (v1.2.1) in R ^53^. In this analysis, first two and last two amino acids of each CDR3b segment were omitted. The TCR logos representing position specific amino acid composition of the CDR3 segments were generated using ggseqlogo (v0.1)^54^.

### CARLIN barcode data processing

Raw sequencing reads were processed using a modified version of our previous 10X single-cell lineage pipeline ^55,56^. Briefly, pair-end reads are combined, and the resulting sequences are collapsed on the 10X cell identifier and unique molecular identifier (UMI). The resulting sequences are aligned to the lineage barcode map (CARLIN array) using the Needleman-Wunsch algorithm with custom parameters detailed in the codebase. The editing outcomes are then called at each target site into a summary file, which includes filters for poor alignment and sequencing primer mismatches. The resulting cell IDs and their CRISPR editing events can then be matched to corresponding entries from the single-cell sequencing data pipelines. The modified pipeline is available on GitHub: https://github.com/mckennalab/SingleCellLineage.

### Constructing CARLIN Clonal Lineage Trees

The *SingleCellLineage* output above was processed using custom scripts to achieve an allele table containing one editing pattern per cell. Trees were built from the allele table using Cassiopeia (v2.0.0), a tool for tree reconstruction for CRISPR/Cas9-based lineage tracing experiments ^57^.

More specifically, the *SingleCellLineage* summary files containing reads collapsed by UMI and corresponding event calls per target site in the CARLIN array were filtered to keep recovered barcodes with two or more UMIs. This threshold is based on Cassiopeia recommendations for cutoffs when working with lineage data. The most frequent allele was used to resolve the cells with multiple alleles. Finally, cells where sequencing reads that did not cover all target sites in the CARLIN array were removed. The lineage groups used in Cassiopeia to identify clones were then defined by a cell’s paired TCRa-TCRb amino acid sequence. Clones with more than three cells were kept, and leaves were annotated by their Seurat cluster.

To obtain clonal lineage trees, the Cassiopeia *VanillaGreedySolver* method was used with default parameters ^57^. Briefly, the Cassiopeia-Greedy algorithm aims to solve the maximum parsimony problem through splitting cells based on the presence or absence of the most frequent mutation. Trees were plotted using the *plot_matplotlib* method from Cassiopeia (matplotlib v3.5.3). Custom pie charts were added to describe the cell-state proportions at internal nodes.

The mega trees in Figure 5 were created with custom javascript code that unified all TCR clones, onto a single figure. Each node contains a pie chart showing the cell-type proportion of all its daughter cells, and terminal nodes are colored by their cell identity (as determined by Seurat) with the CARLIN array editing pattern shown on the outer edge.

## Statistical analysis

Statistical analyses in Figure 3 and Figure 4 were performed with GraphPad Prism 9 (v9.1.2). Statistical analysis was done using one-way ANOVA and Tukey’s multiple comparisons test. P value <0.05 was considered significant.

## Data and Code Availability

All raw and processed single-cell sequencing data (scRNA-seq, scCARLIN-barcode-seq, LR-scTCR-seq) has been deposited at GEO and are publicly available as of the date of publication (GSE241403). All original and modified code used in this study has been deposited at GitHub and is publicly available as of the date of publication. The pipeline used for CARLIN barcode data processing is available at https://github.com/mckennalab/SingleCellLineage. The pipeline used for LR-scTCR-seq is available at https://github.com/indianewok/angerseq. Code written in R and python to process transcriptional data and generate figures 1-4 is available at https://github.com/Leena-Abdullah/CARLIN_CD8_Day_7. Any additional information required to reanalyze the data reported in this paper is available from the lead contact upon request.

**Table S1:** Gene signatures of the CD8 T-cell clusters as determined by Seurat. Related to Figure 1.

**Table S2:** TCR clones in the biased categories with their ES per cluster, frequency, VJ usage and physiochemical properties listed. Related to Figure 3 and Figure 4.

