## Supplemental Data for "Hierarchal single-cell lineage tracing reveals differential fate commitment of CD8 T-cell clones in response to acute infection"

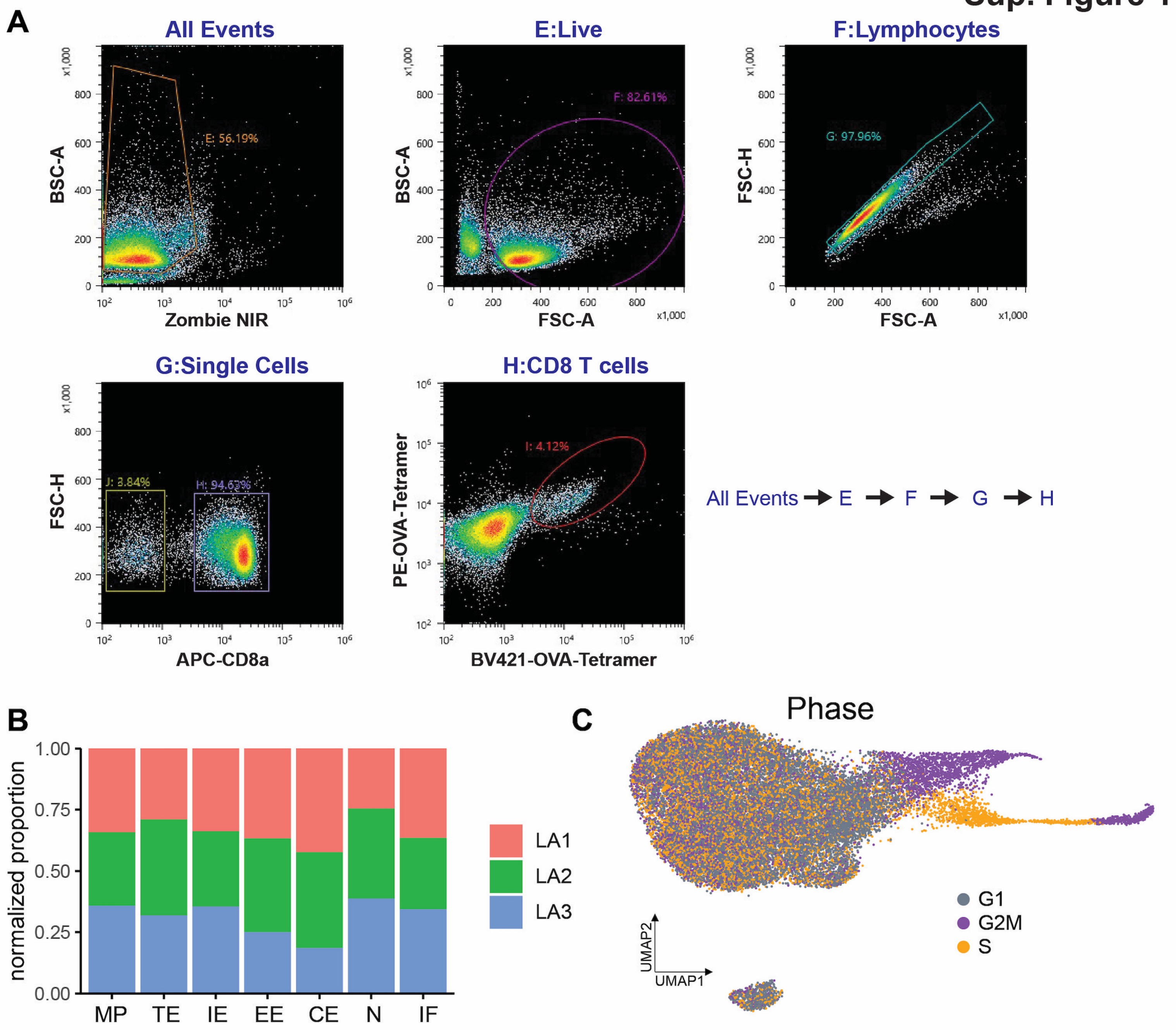


**Figure S1. Flouorescence-activated cell sorting gating strategy along with Seurat cluster composition and cell cycle phase UMAP plot, Related to Figure 1. A.** Flow cytometry gating strategy to sort live, OVA-tetramer+ CD8 T cells from the spleens of infected mice. **B.** Distribution of the 3 samples in the Seurat clusters normalized by sample size. **C.** UMAP plot of cell cycle phase as determined by Seurat.

### Sup. Figure 2

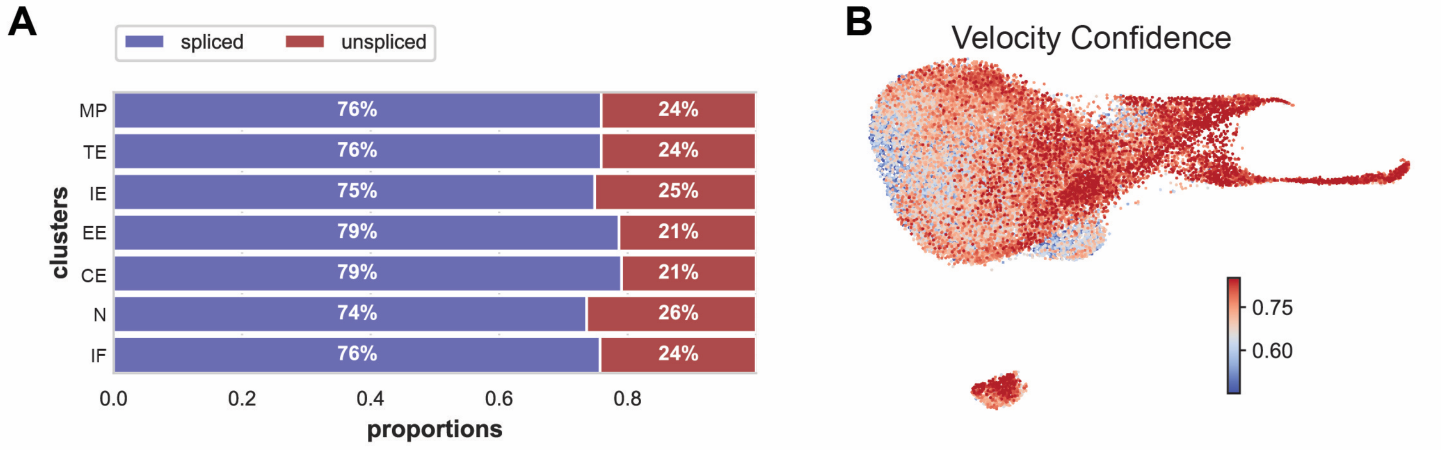


**Figure S2. Percentage of spliced and unspliced counts across clusters and UMAP plot of velocity confidence as determined by scVelo, Related to Figure 2. A.** Proportion of spliced and unspliced counts in each cluster. **B.** Velocity confidence for each cell displayed on the UMAP embedding. Velocity confidence measures coherence between neighboring velocities determined by scVelo.

### Sup. Figure 3

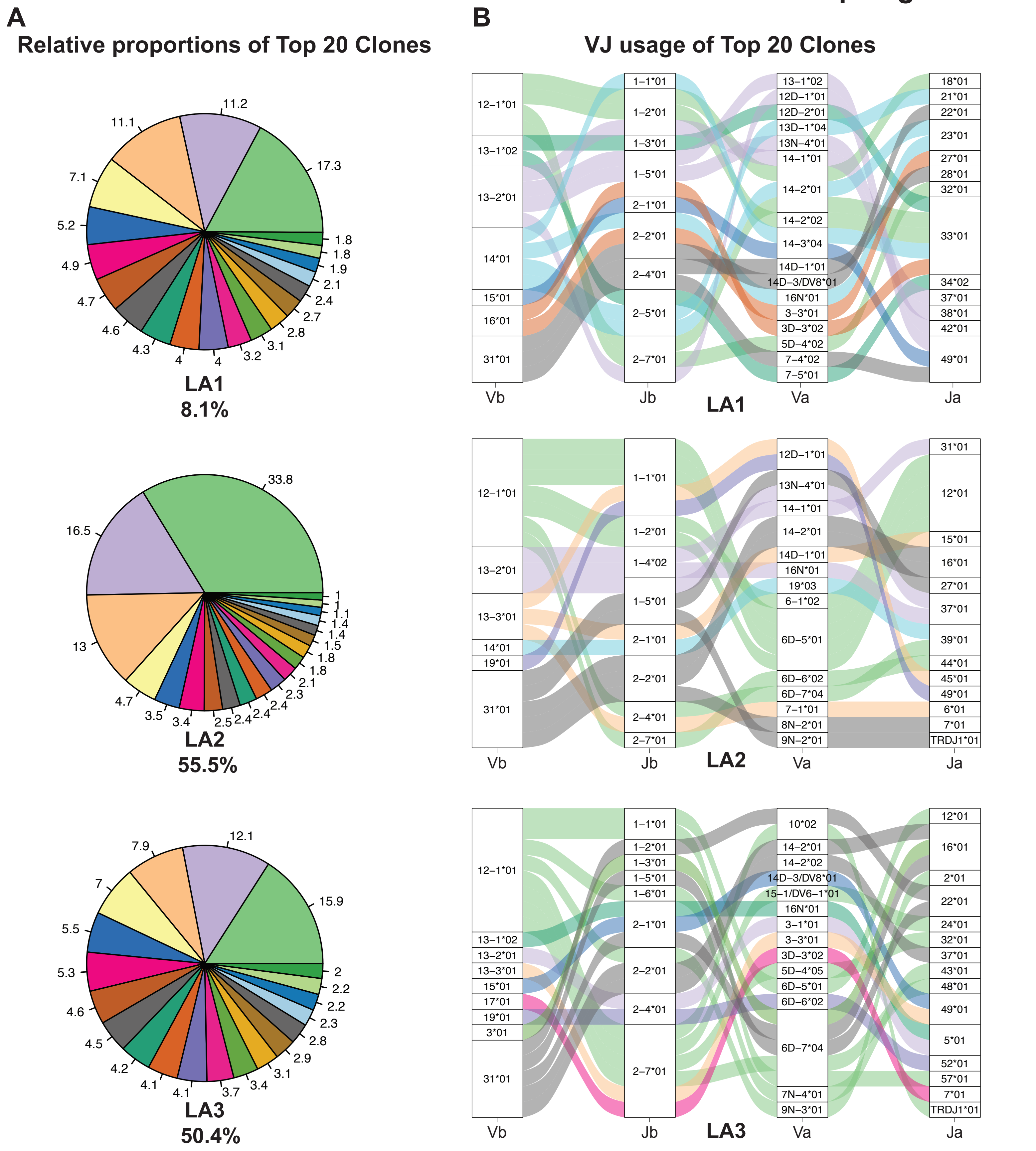


**Figure S3. Relative size and VJ usage of the top 20 clones, Related to Figure 3. A.** Relative proportions of the top 20 clones in the 3 samples. The percentage at the base of each pie chart is the proportion of each sample occupied by the top 20 clones. **B.** Alluvial plots showing the paired VJ alpha and beta usage in the top 20 clones per sample. The stacked bar plots represent the proportion of the TCR segments present in the top 20 clones. The flow streams connecting the stacked bar plots represent the V-J alpha beta usage for the clones. The color of the streams corresponds to the individual Vb segments. Clonal size is not represented in this analysis.

### Sup. Figure 4

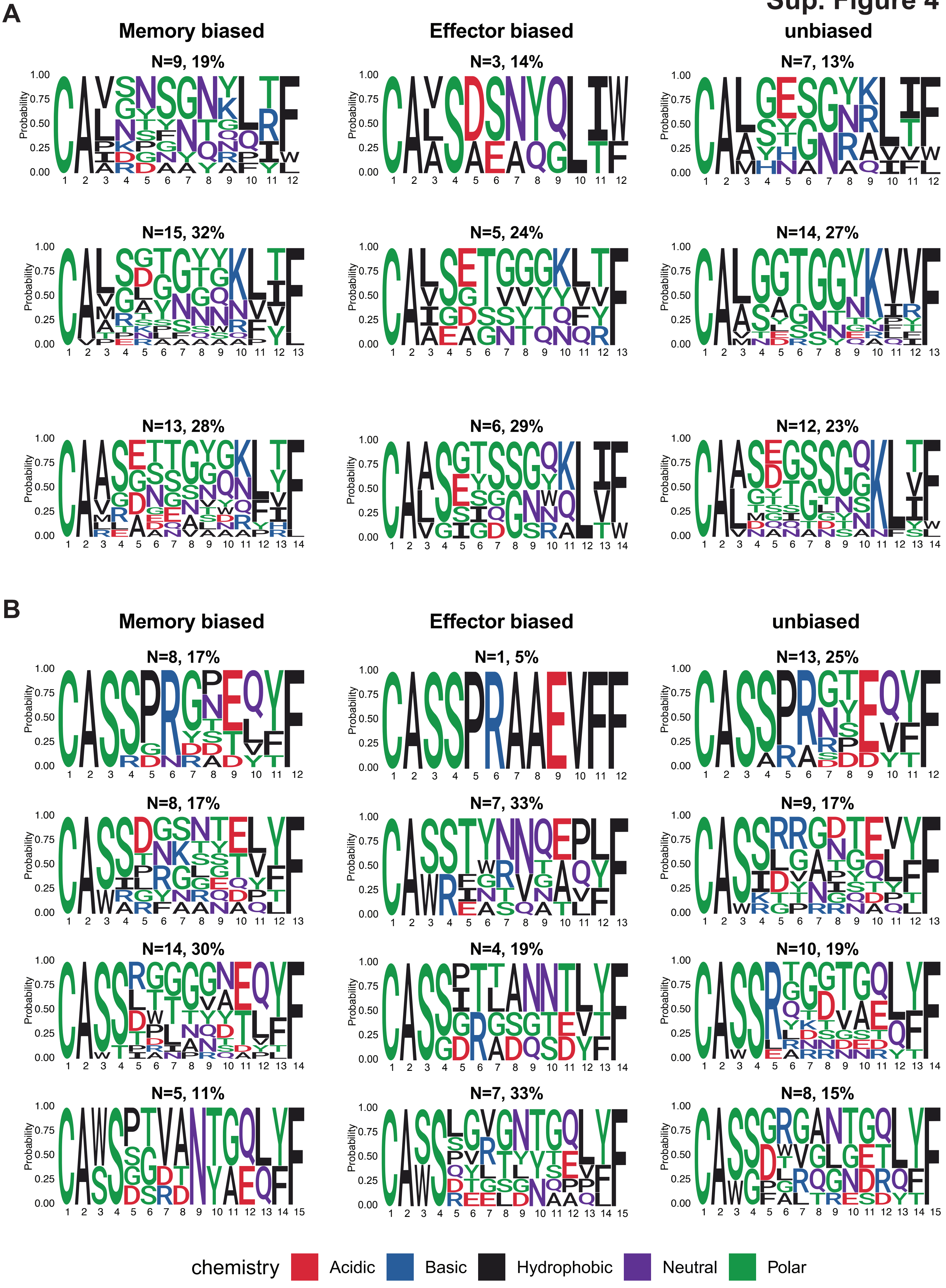

A
Memory biased
Effector biased
unbiased
N=9, 19%
N=3, 14%
N=7, 13%
N=15, 32%
N=5, 24%
N=14, 27%
N=13, 28%
N=6, 29%
N=12, 23%
B
Memory biased
Effector biased
unbiased
N=8, 17%
N=1, 5%
N=13, 25%
N=8, 17%
N=7, 33%
N=9, 17%
N=14, 30%
N=4, 19%
N=10, 19%
N=5, 11%
N=7, 33%
N=8, 15%
Probability
chemistry
Acidic
Basic
Hydrophobic
Neutral
Polar

**Figure S4. CDR3a and CDR3b logos of the biased clones, Related to Figure 4.** Position specific amino acid composition of **A.** CDR3a and **B.** CDR3b segments in the biased clones. Every row represents a specific CDR3 length. The height of the letters represents their frequency at that position. The color of the letters represents the chemical property of the amino acids. Above the TCR logos, N represents the number and the percentage represents the proportion of the biased group represented by the logo.
